# A Systems Neuroscience Approach Identifies IL1B-CASP3 Signaling as a Molecular Link Between Polystyrene Exposure and Alzheimer’s Disease

**DOI:** 10.64898/2026.08.17.745375

**Authors:** Rohan Gupta, Sorabh Lakhanpal, Saurabh Gupta, Sunil Kumar

**Author notes:** Author to whom correspondence should be addressed: Dr. Rohan Gupta, PhD, Assistant Professor, Department of Biotechnology and Bioengineering, School of Biosciences and Technology, Galgotias University, Greater Noida, Uttar Pradesh 203201.

## Abstract

The widespread presence of microplastics and nanoplastics has emerged as a significant environmental concern, with increasing evidence suggesting potential adverse effects on neurological health. However, the molecular mechanisms linking polystyrene exposure to Alzheimer’s disease (AD) remain poorly understood. In this study, an integrative systems biology framework was employed to investigate the molecular interplay between environmental polystyrene exposure and AD pathogenesis. AD-associated genes were retrieved from the Comparative Toxicogenomics Database (CTD) and DisGeNET, while polystyrene-responsive genes were obtained from CTD. Integration of these datasets identified 16 shared genes potentially connecting polystyrene exposure with AD. Transcriptomic analysis of the hippocampal dataset GSE29378 revealed significant differential expression of several overlapping genes between AD and healthy controls. Functional enrichment analyses demonstrated that these genes are predominantly involved in oxidative stress, inflammatory signaling, apoptosis, and synaptic function, all of which are central to AD pathology. Weighted gene co-expression network analysis (WGCNA) further identified disease-associated modules containing multiple intersecting genes strongly correlated with AD clinical traits. Protein–protein interaction analysis highlighted IL1B, CASP3, BCL2, ACHE, and APOE as key hub genes, indicating their potential roles in integrating environmental stress responses with neurodegenerative pathways. Independent validation using the GSE48350 dataset confirmed the robust diagnostic performance of several hub genes in discriminating AD from control samples. Collectively, these findings suggest that environmental polystyrene exposure may promote AD progression through neuroinflammation, oxidative stress, apoptosis, and synaptic dysfunction, providing novel mechanistic insights and identifying promising molecular targets for future experimental, clinical, and epidemiological investigations.

## 1. Introduction

The widespread production and disposal of plastics over the past several decades have resulted in the pervasive presence of microplastics and nanoplastics in the global environment, raising significant concerns regarding their potential impacts on ecological and human health. Microplastics, typically defined as plastic particles smaller than 5 mm, are generated through the degradation of larger plastic materials or directly released from industrial products and consumer goods (Thompson et al., 2004, 2024). Among these, polystyrene, a commonly used synthetic polymer found in packaging materials, disposable containers, and insulation products, represents one of the most prevalent forms of environmental plastic pollution. Due to their small size and high persistence, microplastics can enter biological systems through ingestion, inhalation, or dermal contact, ultimately accumulating in various tissues (Nawab et al., 2024; Kong et al., 2025). Recent studies have demonstrated that microplastics and nanoplastics are capable of crossing biological barriers, including the blood-brain barrier (BBB), thereby raising concerns about their potential neurotoxic effects. In parallel with the growing environmental burden of microplastics, Alzheimer’s disease (AD) has emerged as one of the most pressing global public health challenges (Gou et al., 2024; Nihart et al., 2025; Sun & Song, 2025). AD is a progressive neurodegenerative diseases (NDDs) characterized by cognitive decline, memory impairment, synaptic dysfunction, and neuronal loss, primarily affecting elderly populations. The pathological hallmarks of AD include the accumulation of amyloid-beta plaques, neurofibrillary tangles composed of hyperphosphorylated tau protein, chronic neuroinflammation, mitochondrial dysfunction, and oxidative stress. Although genetic factors, such as mutations in *APP*, *PSEN1*, and *PSEN2* genes contribute to familial forms of AD, the majority of cases are sporadic and are believed to result from a complex interplay between genetic susceptibility and environmental exposures (Zhang et al., 2024a; Stancheva et al., 2025). Increasing evidence suggests that environmental toxicants, including heavy metals, air pollutants, and industrial chemicals, may influence NDDs by promoting oxidative stress, inflammatory responses, and neuronal damage. However, the potential contribution of environmental microplastics to NDDs remains poorly understood (Suman et al., 2024; Lafram et al., 2025a). Emerging experimental evidence indicates that microplastic particles can induce a range of cellular stress responses, including oxidative stress, mitochondrial dysfunction, inflammatory activation, and apoptotic signaling. These biological effects are particularly relevant to AD pathogenesis, as oxidative damage and chronic neuroinflammation are considered central drivers of neuronal degeneration (Gecegelen et al., 2025; Shi et al., 2025; Saha et al., 2026). For instance, exposure to polystyrene microplastics has been reported to induce reactive oxygen species (ROS) production, lipid peroxidation, and disruption of mitochondrial membrane potential in neuronal and non-neuronal cells. Furthermore, microplastics may act as carriers for environmental pollutants and toxic additives, thereby amplifying their biological effects. These findings raise the possibility that chronic exposure to microplastic particles may contribute to NDDs progression by exacerbating molecular pathways associated with neuronal stress and inflammation (Kadac-Czapska et al., 2024; Araújo et al., 2025).

Despite these emerging concerns, the molecular mechanisms linking microplastic exposure with AD-related pathological pathways remain largely unexplored. In particular, there is limited understanding of how environmental polymers, such as polystyrene may influence gene regulatory networks and signaling pathways involved in neurodegeneration (Wang et al., 2024; Siu et al., 2026). Moreover, advances in high-throughput transcriptomic technologies and bioinformatics have enabled researchers to systematically investigate complex biological systems and identify disease-associated molecular signatures. Integrative computational approaches, including gene-disease association databases, transcriptomic profiling, gene co-expression network analysis, and protein-protein interaction (PPI) network modeling, provide powerful tools for uncovering molecular interactions between environmental exposures and disease-related pathways (Zeng & Guo, 2025; Gao et al., 2026). Such approaches can help identify key regulatory genes and signaling networks that may serve as potential biomarkers or therapeutic targets. Among these computational strategies, Weighted Gene Co-expression Network Analysis (WGCNA) has emerged as a robust method for identifying gene modules associated with specific phenotypic traits by analyzing patterns of gene co-expression across transcriptomic datasets. This approach enables the identification of biologically relevant gene clusters and hub genes that play central roles in disease-related molecular networks (Karami et al., 2021; Wang et al., 2023). Similarly, PPI network analysis can provide insights into functional relationships between genes and help identify critical regulatory nodes within complex biological systems. When combined with functional enrichment analyses, such as Gene Ontology (GO), Kyoto Encyclopedia of Genes and Genomes (KEGG), and Gene Set Enrichment Analysis (GSEA), these methods can reveal the biological processes and signaling pathways underlying disease mechanisms.

Given the increasing environmental prevalence of microplastics and the growing burden of NDDs, there is an urgent need to better understand the potential molecular links between environmental plastic exposure and AD pathogenesis. Identifying such links could provide valuable insights into environmental risk factors contributing to neurodegeneration and may facilitate the development of novel preventive and therapeutic strategies (Yu et al., 2025b). In this context, integrative bioinformatics approaches offer an effective strategy for exploring complex gene–environment interactions by integrating diverse biological datasets. Therefore, the present study aimed to systematically investigate the potential molecular relationship between polystyrene exposure and AD using an integrative systems biology framework. AD-associated genes were collected from curated databases including the Comparative Toxicogenomics Database (CTD) and DisGeNET, while polystyrene-associated genes were obtained from CTD. These datasets were integrated to identify overlapping genes potentially linking environmental plastic exposure with AD-related molecular pathways. Furthermore, transcriptomic analysis was performed using the GSE29378 dataset derived from hippocampal tissues, followed by differential gene expression analysis, gene co-expression network construction using WGCNA, functional enrichment analysis, and PPI network modeling. Finally, key hub genes were validated using an independent dataset to evaluate their diagnostic relevance. By integrating environmental exposure data with transcriptomic and network-based analyses, this study seeks to identify critical genes and signaling pathways that may mediate the interaction between polystyrene microplastic exposure and AD pathogenesis. The findings of this study may provide new insights into the potential role of environmental plastic pollution in NDDs development and highlight candidate molecular targets for future experimental and epidemiological investigations.

## 2. Methodology

### 2.1. Identification and integrative analysis of Alzheimer’s disease- and polystyrene-associated genes

AD-related genes were systematically collected from the CTD (https://ctdbase.org/) and DisGeNET (https://disgenet.com/), two widely used resources that curate and integrate gene-disease associations from experimental studies and the literature. In CTD, genes related to the disease term “Alzheimer Disease” were retrieved, including both curated and inferred associations. In DisGeNET, genes linked to the corresponding disease concept were extracted and filtered to retain associations with a DisGeNET score ≥ 0.1 to ensure moderate to high confidence. Parallelly, polystyrene-related genes were obtained from the CTD by querying the chemical term “Polystyrene”. Genes documented to interact with polystyrene or polystyrene-derived micro/nanoplastics through expression regulation or functional modulation were selected. Only associations supported by curated evidence or experimental reports were included, while entries based solely on predictive inference were excluded. The AD-related gene list was formed by merging genes from CTD and DisGeNET, and the list of genes associated with polystyrene consisted of genes retrieved from CTD. The integration of two gene sets was performed by Venn diagram analysis to identify potential molecular links between polystyrene exposure and AD. Overlapping genes were determined and visualized using the online tool E Venn 2.0. (https://www.bic.ac.cn/EVenn/), which also allowed the extraction of the intersecting gene list for further analyses. Genes that appeared in at least one AD-related database and were simultaneously associated with polystyrene in CTD were considered candidate genes **(Fig. 1) (Supplementary Table 1)**.

**Figure 1:**
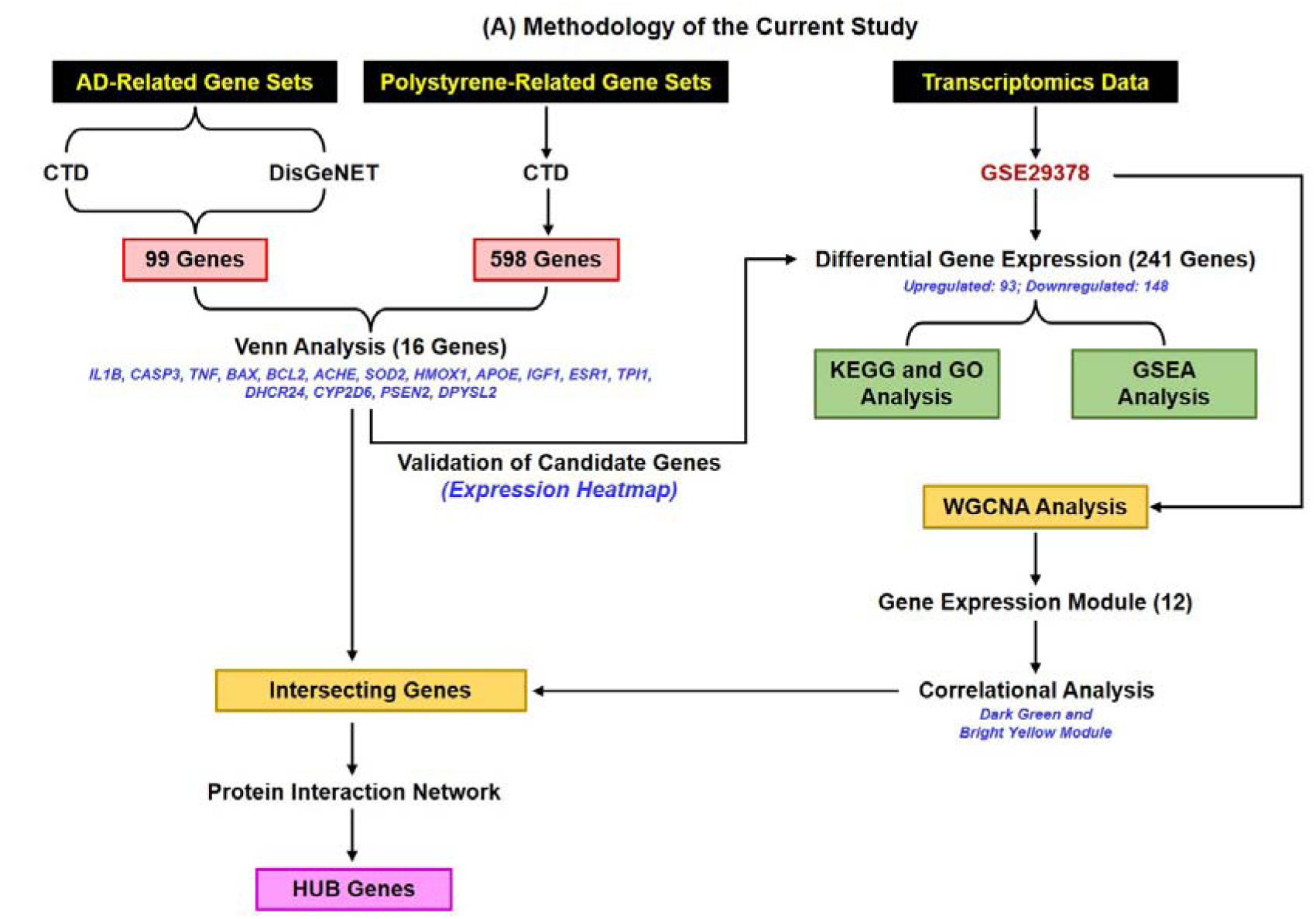
Workflow illustrating the integrative systems biology strategy used to identify molecular links between polystyrene exposure and Alzheimer’s disease. Alzheimer’s disease-related genes were retrieved from the Comparative Toxicogenomics Database and DisGeNET, yielding 99 overlapping genes, while 598 polystyrene-associated genes were obtained from CTD. Venn analysis identified 16 intersecting genes shared between polystyrene exposure and Alzheimer’s disease. Transcriptomic data from the GEO dataset GSE29378 were analyzed to identify differentially expressed genes, followed by GO, KEGG, and GSEA enrichment analyses. WGCNA identified 12 co-expression modules, of which the dark green and bright yellow modules showed the strongest association with Alzheimer’s disease. Intersecting genes present within disease-associated modules were subsequently integrated into a protein–protein interaction network to identify central hub genes.

### 2.2. Transcriptomics analysis of gene expression datasets

Data on gene expression were retrieved from the Gene Expression Omnibus (GEO) (https://www.ncbi.nlm.nih.gov/geo/), a public repository where high-throughput functional genomic datasets are archived, including microarray and other sequencing-based expression profiles. AD-related dataset, GSE29378 (Miller et al., 2013), were downloaded from GEO. In GSE29378, expression profiles derived from hippocampal tissue were extracted and subsequently divided into AD and non-demented control groups. Raw expression data were preprocessed and normalized across the samples, with subsequent log transformation. We then used the limma package in R (version 4.1.3) to identify genes differentially expressed between AD and control groups. For a given gene, |log FC| > 0.5 and nominal p < 0.05 were considered statistically significant. To enhance the estimation of variance, empirical Bayes moderation was performed with the application of limma as proposed by Ritchie et al., 2015 (Ritchie et al., 2015). All statistical analyses were performed in R, and p < 0.05 was considered indicative of statistical significance. Data visualization was performed using the ggplot2 package (https://github.com/tidyverse/ggplot2), and heat maps were created using the pheatmap package (https://cran.r-project.org/web/packages/pheatmap/index.html). The identified differential expressed genes (DEGs) were further visualized on volcano plots, heatmaps, and PCA plots to depict their expression pattern and overall distribution.

### 2.3. WGCNA network construction and identification of AD-related modules

Weighted gene co-expression network analysis was performed using the R package WGCNA to build a co-expression network from the gene expression dataset GSE29378 according to Langfelder and Horvath (2008) (Langfelder & Horvath, 2008). To reduce noise and computational burden, MAD was calculated for each gene, and then the half of genes with the lowest variability were excluded. The outlier genes and samples were further removed by the goodSamplesGenes function in the WGCNA package. Remaining genes were used for building a scale-free co-expression network. Moreover, Pearson correlation coefficients were calculated on a pairwise basis to produce a similarity matrix, and the latter was then thresholded into an adjacency matrix using soft-thresholding with power β to emphasize strong correlations while giving lower weights to weaker correlations. Scale-free topology criteria were followed to set β to 16. The adjacency matrix was then converted to a Topological Overlap Matrix (TOM), and the TOM-based dissimilarity measure, 1 − TOM, was computed to estimate network connectivity. Genes were clustered hierarchically using TOM dissimilarity, in which the minimum module size was defined as 30 genes and a deepSplit (sensitivity) parameter of 3. When a cut height of 0.25 was used, similar modules were merged into 12 co-expression modules; those genes which could not be assigned to any module were placed into the gray module **(Fig. 1)**.

### 2.4. Module-trait relationship analysis and hub gene extraction

The eigengenes of modules were then correlated with clinical traits, such as AD status, sex, and hippocampal subregions, by using Pearson correlation analysis. Modules in which the correlation of eigengenes with AD-related traits was significant (p < 0.05) were selected. For each gene, module membership (MM) and gene significance for AD (GS) were calculated to extract biologically relevant genes. Modules which showed the strongest relationship with AD, especially bright yellow and dark green modules, were selected, and the genes belonging to these modules were retrieved as candidate hub genes. GO (https://geneontology.org/docs/go-enrichment-analysis/) and KEGG pathway (https://www.genome.jp/kegg/pathway.html) enrichment analyses were performed for functional annotation of these genes to understand the biological processes and pathways in AD pathogenesis.

### 2.5. PPI Network Construction and Diagnostic Hub Gene Validation

To elucidate the functional interplay between the identified intersecting genes, a PPI network was constructed using the STRING database (https://string-db.org/) with a composite confidence score threshold of 0.4. The resulting interactome was visualized in Cytoscape (https://cytoscape.org/), where the CytoHubba plugin (Chin et al., 2014) was employed to identify the top ten hub genes based on topological network analysis. These hub genes were subsequently validated for their diagnostic relevance to AD using transcriptomic data from the GSE48350 dataset, specifically comparing normal and AD-affected hippocampal tissues. The diagnostic efficacy of these candidates was quantified via Receiver Operating Characteristic (ROC) curve analysis utilizing the “pROC” R package (https://xrobin.github.io/pROC/); genes demonstrating an Area Under the Curve (AUC) exceeding 0.6 were identified as diagnostic markers possessing a statistically relevant degree of predictive accuracy **(Fig. 1)**.

## 3. Results

### 3.1. Integrative CTD-DisGeNET Analysis Identifies Shared Genes Linking Polystyrene Exposure to Alzheimer’s Disease

To investigate potential molecular interactions between AD and polystyrene exposure, an integrative database mining approach was performed using the CTD and DisGeNET, two widely recognized repositories of curated gene-disease associations. Initially, AD-related genes were retrieved independently from both databases and subsequently integrated to obtain a consolidated gene set. Further, venn diagram analysis revealed 99 overlapping genes shared between CTD and DisGeNET **(Supplementary Table 2)**, indicating a substantial consensus between curated experimental evidence and literature-derived associations **(Fig. 2A)**. Specifically, 11 genes were uniquely identified in CTD, while 112 genes were uniquely identified in DisGeNET, reflecting differences in database coverage and annotation strategies. Further, to explore the potential impact of environmental plastic exposure on AD-related molecular pathways, genes associated with polystyrene were retrieved from CTD and intersected with the AD gene set. Integration of these datasets revealed 16 shared genes that were simultaneously associated with AD and polystyrene exposure **(Fig. 2A) (Supplementary Table 2)**. These genes included IL1B, CASP3, TNF, BAX, BCL2, ACHE, SOD2, HMOX1, APOE, IGF1, ESR1, TPI1, DHCR24, CYP2D6, PSEN2, and DPYSL2. Functional annotation of these genes suggests their involvement in several core biological processes implicated in neurodegeneration, including neuroinflammation, mitochondrial dysfunction, oxidative stress, apoptotic signaling, and neuronal metabolism. The identification of these overlapping genes therefore provides a preliminary molecular framework linking environmental microplastic exposure to AD-related pathological mechanisms.

**Figure 2:**
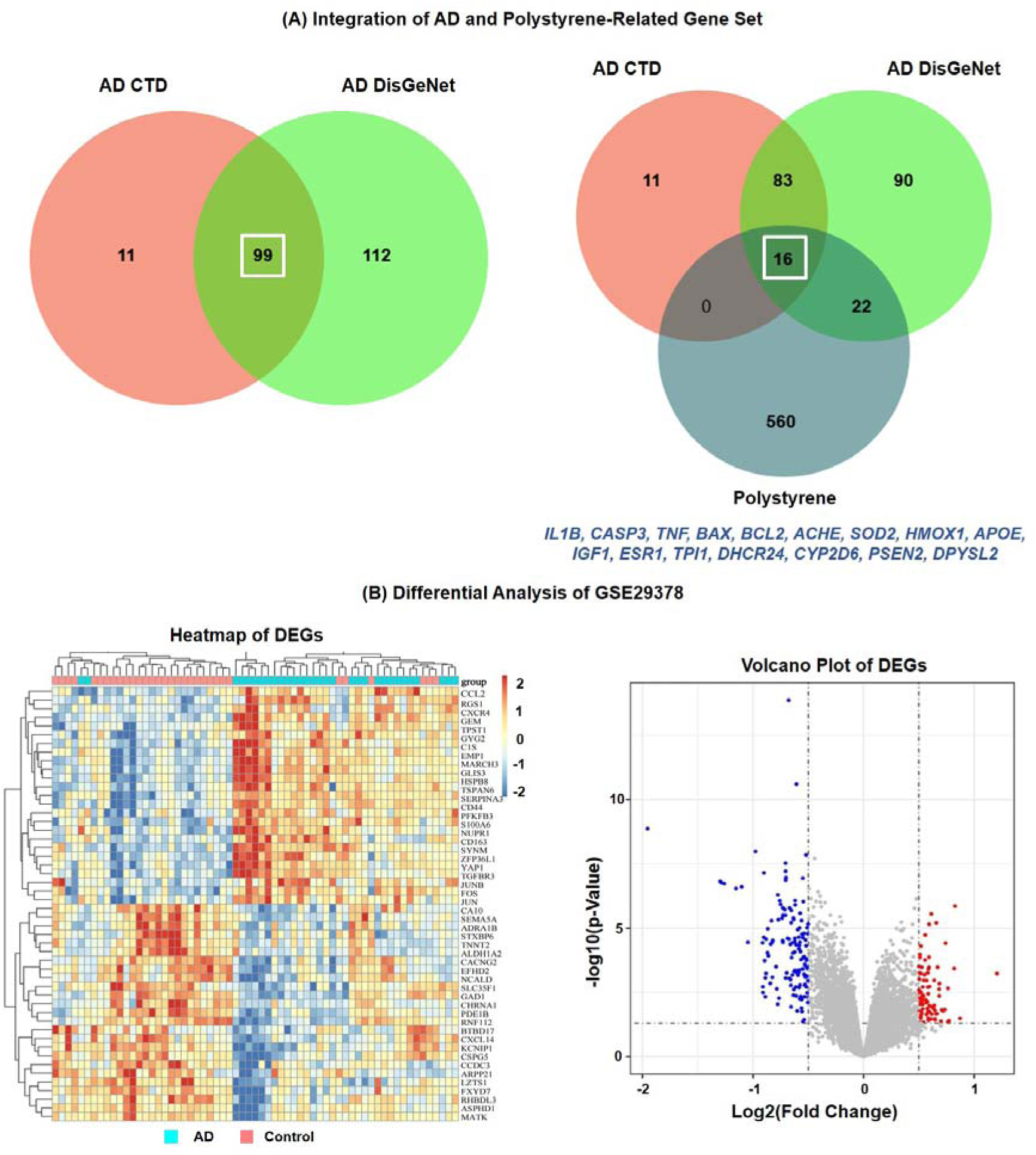
Identification of genes linking polystyrene exposure with Alzheimer’s disease and transcriptomic alterations in hippocampal tissue. (A) Venn diagram showing overlap between Alzheimer’s disease-related genes retrieved from the Comparative Toxicogenomics Database and DisGeNET. A total of 99 genes were common between the two databases. Integration of these genes with polystyrene-associated genes identified 16 intersecting genes shared among AD CTD, AD DisGeNET, and polystyrene datasets: IL1B, CASP3, TNF, BAX, BCL2, ACHE, SOD2, HMOX1, APOE, IGF1, ESR1, TPI1, DHCR24, CYP2D6, PSEN2, and DPYSL2. (B) Differential expression analysis of the GSE29378 hippocampal dataset. The heatmap illustrates the expression patterns of differentially expressed genes between AD and control samples, showing clear clustering of the two groups. The volcano plot displays significantly upregulated genes in red and downregulated genes in blue based on |log2FC| > 0.5 and p < 0.05.

### 3.2. Transcriptomic Profiling Reveals Significant Gene Expression Alterations in AD Hippocampal Tissue

To validate the relevance of the identified genes within the AD transcriptomic landscape, gene expression analysis was performed using the GSE29378 dataset, which contains microarray data derived from hippocampal tissues of AD patients and non-demented controls. Following normalization and log transformation, differential gene expression analysis was conducted using the limma algorithm, applying thresholds of |log FC| > 0.5 and p < 0.05 **(Supplementary Table 3)**. Additionally, visualization of the differential expression results through volcano plot analysis revealed widespread transcriptional dysregulation between AD and control samples (Fig. 2B). Numerous genes exhibited statistically significant expression changes, indicating large-scale molecular alterations associated with AD pathology. Hierarchical clustering of DEGs further demonstrated clear separation between AD and control samples, confirming the robustness of the transcriptomic signatures identified in the dataset **(Fig. 2B)**. Among the 16 intersecting genes identified through database integration, several exhibited significant differential expressions within the AD dataset. Specifically, ESR1 (logFC = −1.05), PSEN2 (logFC = −0.71), TNF (logFC = −0.70), APOE (logFC = −0.63), IGF1 (logFC = −0.59), and BCL2 (logFC = −0.57) were significantly downregulated in AD hippocampal tissues. In contrast, genes including BAX (logFC = 1.02), SOD2 (logFC = 0.90), DHCR24 (logFC = 0.88), HMOX1 (logFC = 0.85), CASP3 (logFC = 0.84), and TPI1 (logFC = 0.75) were significantly upregulated **(Table 1) (Supplementary Fig. 1)**. These expression changes collectively suggest activation of pro-apoptotic signaling pathways, oxidative stress responses, and metabolic adaptation processes in AD brains. Consistent with these findings, heatmap visualization of the intersecting genes demonstrated clear expression differences between AD and control samples, revealing two major clusters corresponding to upregulated stress-response genes and downregulated neuroprotective genes **(Fig. 3A)**. These transcriptional alterations highlight the involvement of environmental stress-responsive pathways in AD pathogenesis.

**Figure 3:**
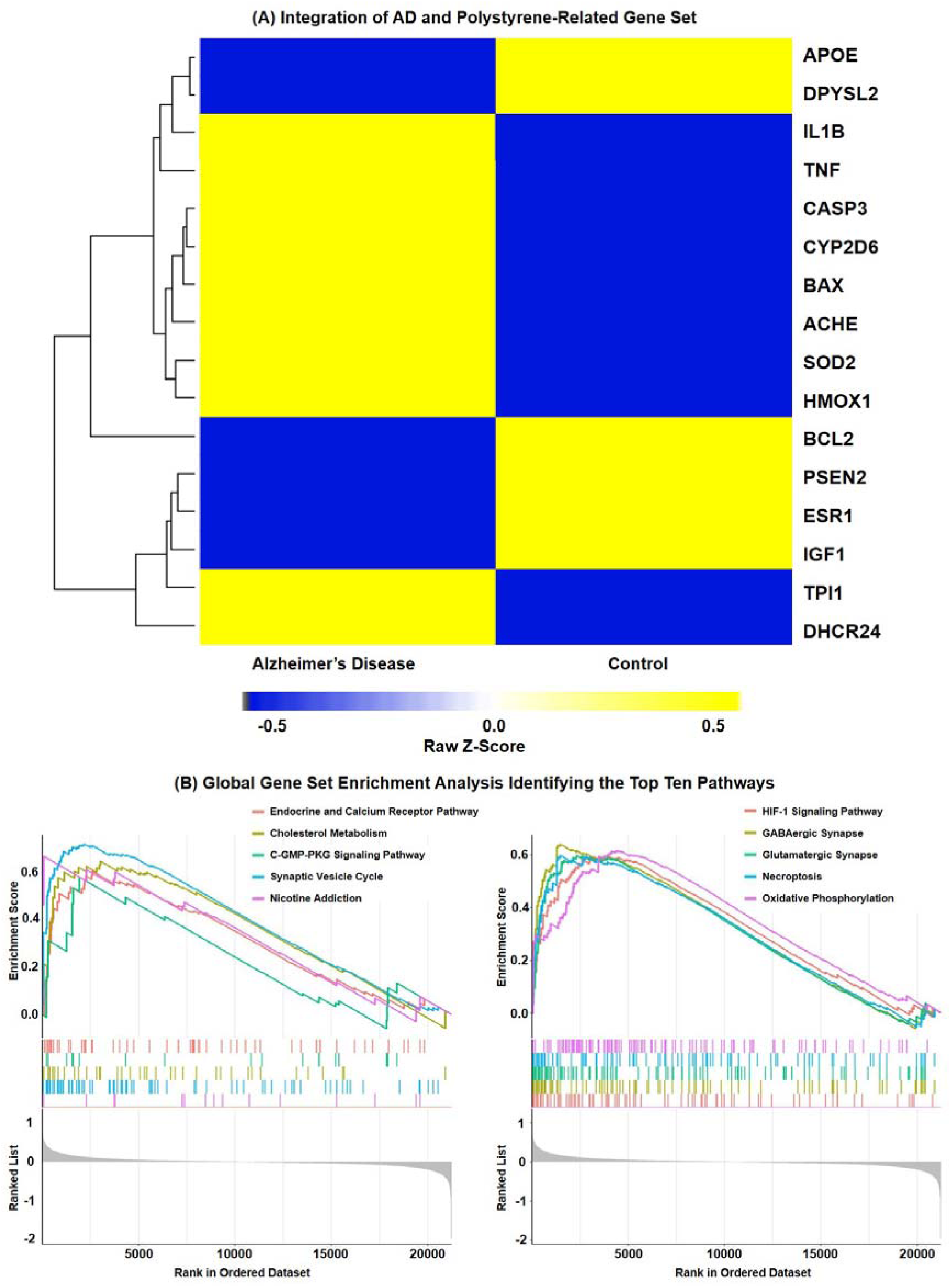
Expression pattern and pathway enrichment analysis of genes linking polystyrene exposure with Alzheimer’s disease. (A) Heatmap showing relative expression of the 16 intersecting genes in Alzheimer’s disease and control hippocampal samples from GSE29378. Yellow indicates higher expression and blue indicates lower expression based on row Z-scores. Genes including IL1B, TNF, CASP3, BAX, ACHE, SOD2, HMOX1, CYP2D6, TPI1, and DHCR24 were upregulated in AD, whereas APOE, DPYSL2, BCL2, PSEN2, ESR1, and IGF1 were downregulated. (B) Gene Set Enrichment Analysis identifying the top enriched pathways in AD hippocampal tissues. Significantly enriched pathways included endocrine and calcium receptor signaling, cholesterol metabolism, cGMP–PKG signaling, synaptic vesicle cycle, nicotine addiction, HIF-1 signaling, GABAergic synapse, glutamatergic synapse, necroptosis, and oxidative phosphorylation.

**Table 1:** Differential expression analysis of the 16 intersecting genes linking polystyrene exposure and Alzheimer’s disease in the GSE29378 hippocampal dataset. The table summarizes adjusted p values, log2 fold change, t-statistics, and gene annotations for candidate genes identified through integrative analysis of polystyrene- and Alzheimer’s disease-associated gene sets. Negative log2FC values indicate downregulation in AD samples, whereas positive values indicate upregulation.

| ID | adj.P.Val | P.Value | t | B | logFC | Gene Symbol | Gene Title |
| --- | --- | --- | --- | --- | --- | --- | --- |
| 10025920920.00 | 0.00 | 0.00 | 5.62 | 7.68 | -1.05 | ESR1 | estrogen receptor 1 |
| 10023819976.00 | 0.00 | 0.00 | -13.39 | 68.24 | -0.71 | PSEN2 | presenilin 2 (Alzheimer disease 4) |
| 10025909848.00 | 0.00 | 0.00 | 4.45 | 2.09 | -0.7 | TNF | tumor necrosis factor (TNF superfamily) |
| 10023843485.00 | 0.78 | 0.00 | -0.32 | -7.54 | -0.65 | DPYSL2 | dihydropyrimidinase-like 2 |
| 10033668702.00 | 0.00 | 0.00 | 7.55 | 19.31 | -0.63 | APOE | apolipoprotein E |
| 10025906631.00 | 0.00 | 0.00 | -20.78 | 145.16 | -0.59 | IGF1 | insulin-like growth factor 1 (somatomedin C) |
| 10023820570.00 | 0.00 | 0.00 | 14.44 | 78.58 | -0.57 | BCL2 | B-cell CLL/lymphoma 2 |
| 10025908937.00 | 0.00 | 0.00 | 4.37 | 1.78 | -0.49 | IL1B | interleukin 1 |
| 10033668469.00 | 0.00 | 0.00 | 7.20 | 16.99 | 0.55 | CYP2D6 | cytochrome P450 |
| 10025908068.00 | 0.31 | 0.00 | -1.11 | -6.98 | 0.56 | ACHE | acetylcholinesterase (Yt blood group) |
| 10023809797.00 | 0.00 | 0.00 | -16.39 | 98.42 | 0.75 | TPI1 | triosephosphate isomerase 1 |
| 10025910100.00 | 0.17 | 0.00 | -1.50 | -6.47 | 0.84 | CASP3 | caspase 3 |
| 10023806944.00 | 0.00 | 0.00 | 14.09 | 75.11 | 0.85 | HMOX1 | heme oxygenase (decycling) 1 |
| 10025903870.00 | 0.00 | 0.00 | -6.32 | 11.56 | 0.88 | DHCR24 | 24-dehydrocholesterol reductase |
| 10023816501.00 | 0.00 | 0.00 | 10.53 | 42.17 | 0.9 | SOD2 | superoxide dismutase 2 |
| 10023812217.00 | 0.00 | 0.00 | 8.64 | 27.01 | 1.02 | BAX | BCL2-associated X protein |

### 3.3. Pathway Enrichment Analysis Reveals Dysregulation of Neuroinflammatory and Synaptic Signaling Pathways

To elucidate the biological pathways associated with the identified transcriptional alterations, GSEA was conducted. The analysis identified several significantly enriched pathways associated with neuronal signaling, metabolic regulation, and neurodegenerative processes **(Fig. 3B)**. Notably, pathways related to synaptic function and neuronal signaling were significantly enriched, including the GABAergic synapse, glutamatergic synapse, and synaptic vesicle cycle pathways, suggesting impaired neurotransmission in AD. Additionally, enrichment of cholesterol metabolism and cGMP-PKG signaling pathways indicates potential disruptions in neuronal lipid homeostasis and intracellular signaling cascades that are critical for neuronal survival and plasticity. Importantly, pathways associated with cell death and oxidative stress, including necroptosis and oxidative phosphorylation, were also enriched. These pathways have been widely implicated in NDDs and are consistent with the hypothesis that environmental toxicants may exacerbate neurodegeneration through oxidative stress and mitochondrial dysfunction. Further pathway enrichment using Reactome analysis revealed significant involvement of immune-related signaling pathways, including interleukin signaling, pyroptosis, inflammasome activation, and programmed cell death pathways **(Supplementary Fig. 2)**. These findings strongly support the notion that neuroinflammatory processes represent a central component of AD pathology and may be modulated by environmental exposures such as microplastics. Additionally, GO analysis further corroborated these observations, highlighting enrichment of biological processes, such as amyloid precursor protein metabolism, regulation of neuronal apoptosis, nitric oxide biosynthesis, and amyloid-beta formation. Collectively, these results suggest that the identified genes participate in interconnected biological processes that converge on neuronal degeneration and inflammatory responses.

### 3.4. Construction of Co-expression Networks Identifies AD-Associated Gene Modules

To further investigate the transcriptional architecture underlying AD pathology, WGCNA was performed using the GSE29378 dataset. To construct a biologically meaningful network, the optimal soft-thresholding power was determined based on scale-free topology criteria. A soft-thresholding power of β = 16 was selected as it achieved a scale-free topology fit index (R² ≈ 0.87) while maintaining sufficient network connectivity **(Fig. 4A)**. Hierarchical clustering based on the TOM identified 12 distinct gene modules, each representing clusters of highly co-expressed genes **(Fig. 4B)**. Genes that did not belong to any module were grouped into the gray module, representing genes with weak or inconsistent correlations. Further, sample clustering analysis revealed clear segregation of AD and control samples, further validating the robustness of the dataset and confirming the presence of disease-associated transcriptional patterns **(Fig. 4C)**. In addition to disease status, the clustering analysis also captured biological variation associated with sex and hippocampal subregions (CA1 and CA3), highlighting the complex transcriptional heterogeneity within the hippocampus.

**Figure 4:**
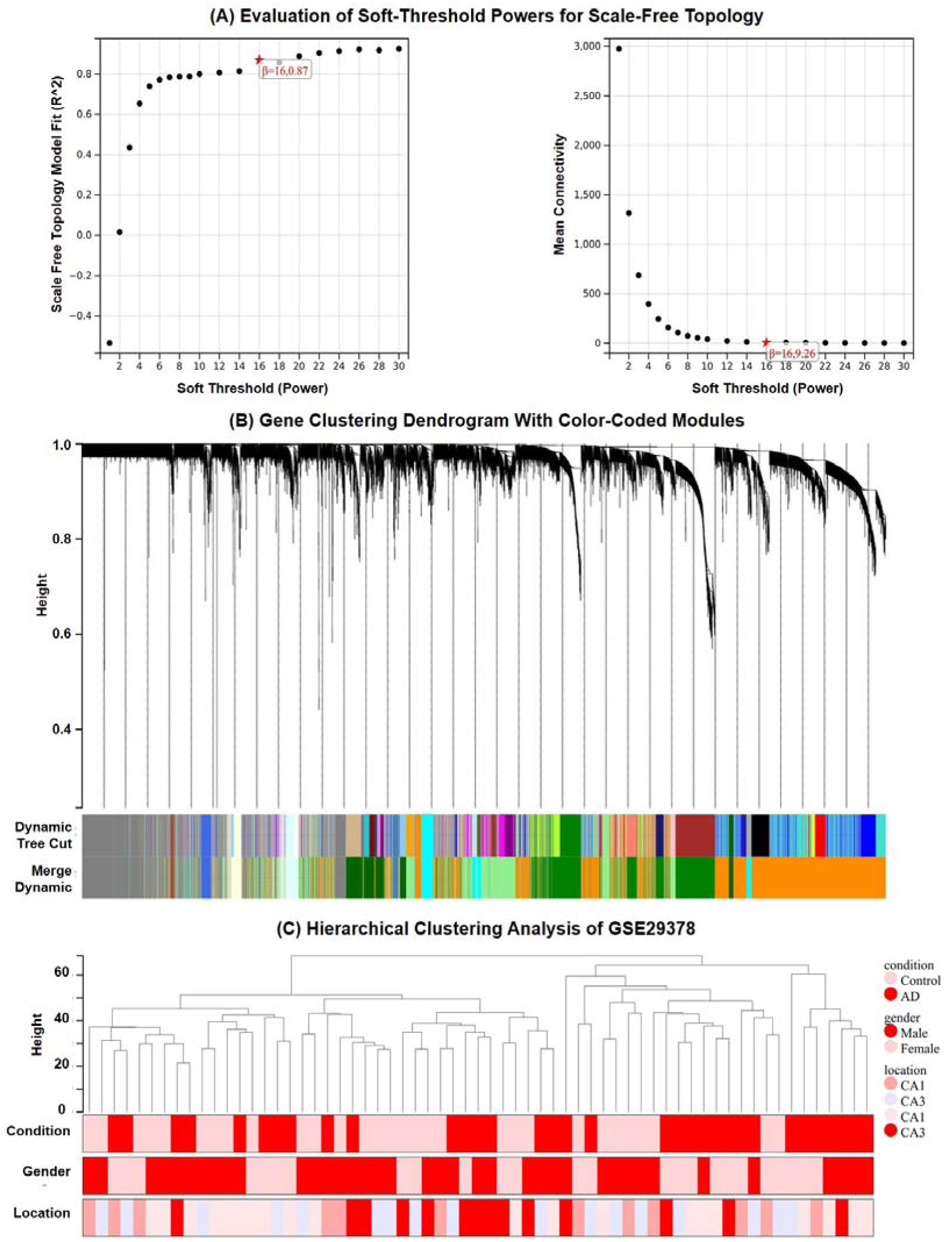
Weighted gene co-expression network analysis of the GSE29378 hippocampal dataset. (A) Determination of the optimal soft-thresholding power for construction of a scale-free network. The left panel shows the scale-free topology fit index (R²) across different soft-threshold powers, while the right panel shows mean connectivity. A soft-thresholding power of β = 16 was selected to achieve an approximate scale-free topology with sufficient network connectivity. (B) Hierarchical clustering dendrogram of genes based on topological overlap, with branches corresponding to co-expression modules identified by dynamic tree cutting. The lower color bars indicate the initial dynamic tree cut modules and the merged modules after module eigengene clustering. (C) Hierarchical clustering of samples from the GSE29378 dataset according to gene expression profiles. Colored annotation bars represent clinical condition (AD or control), sex, and hippocampal subregion (CA1 or CA3), demonstrating clear transcriptional segregation between groups.

### 3.5. Identification of AD-Associated Modules Through Module-Trait Correlation Analysis

To identify gene modules associated with clinical traits, correlations between module eigengenes and phenotypic features were calculated. The module-trait relationship heatmap revealed several modules significantly associated with AD status **(Fig. 5A)**. Among these, the dark green and light-yellow modules exhibited the strongest correlations with AD, suggesting that genes within these modules may play key roles in AD pathogenesis **(Supplementary Table 4 and Supplementary Table 5)**. Integration of the intersecting AD-polystyrene genes with WGCNA modules revealed that seven genes (IL1B, CASP3, TNF, BCL2, ACHE, APOE, and IGF1) were located within the dark green module, while two genes (PSEN2 and DPYSL2) were located within the light-yellow module **(Fig. 5B)**. Moreover, scatter plot analysis of MM and GS further demonstrated strong positive correlations between gene connectivity and disease association, confirming the biological relevance of these modules **(Supplementary Fig. 3)**. These results indicate that the identified modules represent key transcriptional networks involved in neuroinflammation, apoptosis, and neuronal dysfunction in AD.

**Figure 5:**
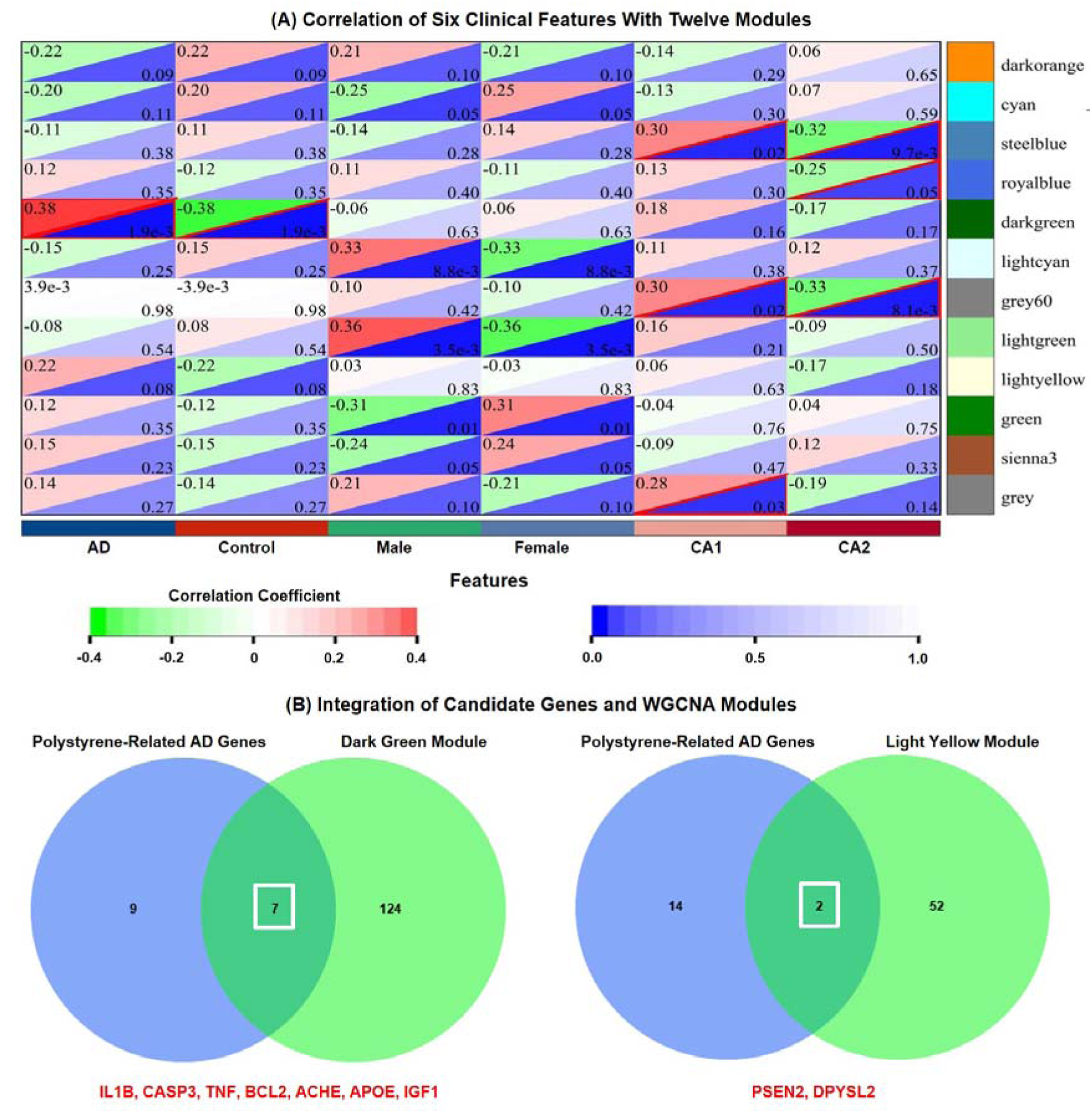
Identification of Alzheimer’s disease-associated co-expression modules and integration with polystyrene-responsive genes. (A) Module–trait relationship heatmap showing correlations between the 12 WGCNA modules and clinical features, including AD status, control status, sex, and hippocampal subregions. Red indicates positive correlations and green indicates negative correlations. The dark green and light-yellow modules exhibited the strongest associations with Alzheimer’s disease. (B) Venn diagram analysis integrating the 16 polystyrene-related Alzheimer’s disease genes with genes from the dark green and light-yellow WGCNA modules. Seven genes (IL1B, CASP3, TNF, BCL2, ACHE, APOE, and IGF1) overlapped with the dark green module, whereas two genes (PSEN2 and DPYSL2) overlapped with the light-yellow module.

### 3.6. Protein-Protein Interaction Network Reveals Central Hub Genes

To further characterize the functional relationships among the candidate genes, a PPI network was constructed using the STRING database and visualized in Cytoscape. The resulting network revealed extensive interactions among genes involved in inflammatory signaling, apoptosis, and neuronal metabolism **(Fig. 6)**. Topological analysis using the CytoHubba algorithm identified BCL2, IL1B, CASP3, ACHE, and APOE as the most highly connected hub genes within the network. These genes represent central regulatory nodes in several biological processes critical to AD pathogenesis. Specifically, IL1B functions as a key mediator of neuroinflammation, CASP3 and BCL2 regulate apoptotic signaling pathways, ACHE modulates cholinergic neurotransmission, and APOE plays a pivotal role in lipid metabolism and amyloid-beta clearance. The convergence of these pathways highlights a potential mechanistic link between environmental stress responses and neurodegenerative signaling cascades. Moreover, to evaluate the potential clinical relevance of the identified hub genes, ROC curve analysis was performed using an independent validation dataset (GSE48350). The analysis demonstrated strong diagnostic performance for several hub genes. Among the evaluated candidates, BCL2 exhibited the highest diagnostic accuracy (AUC = 0.929), followed by IL1B (AUC = 0.898) and CASP3 (AUC = 0.842) **(Fig. 6)**.

**Figure 6:**
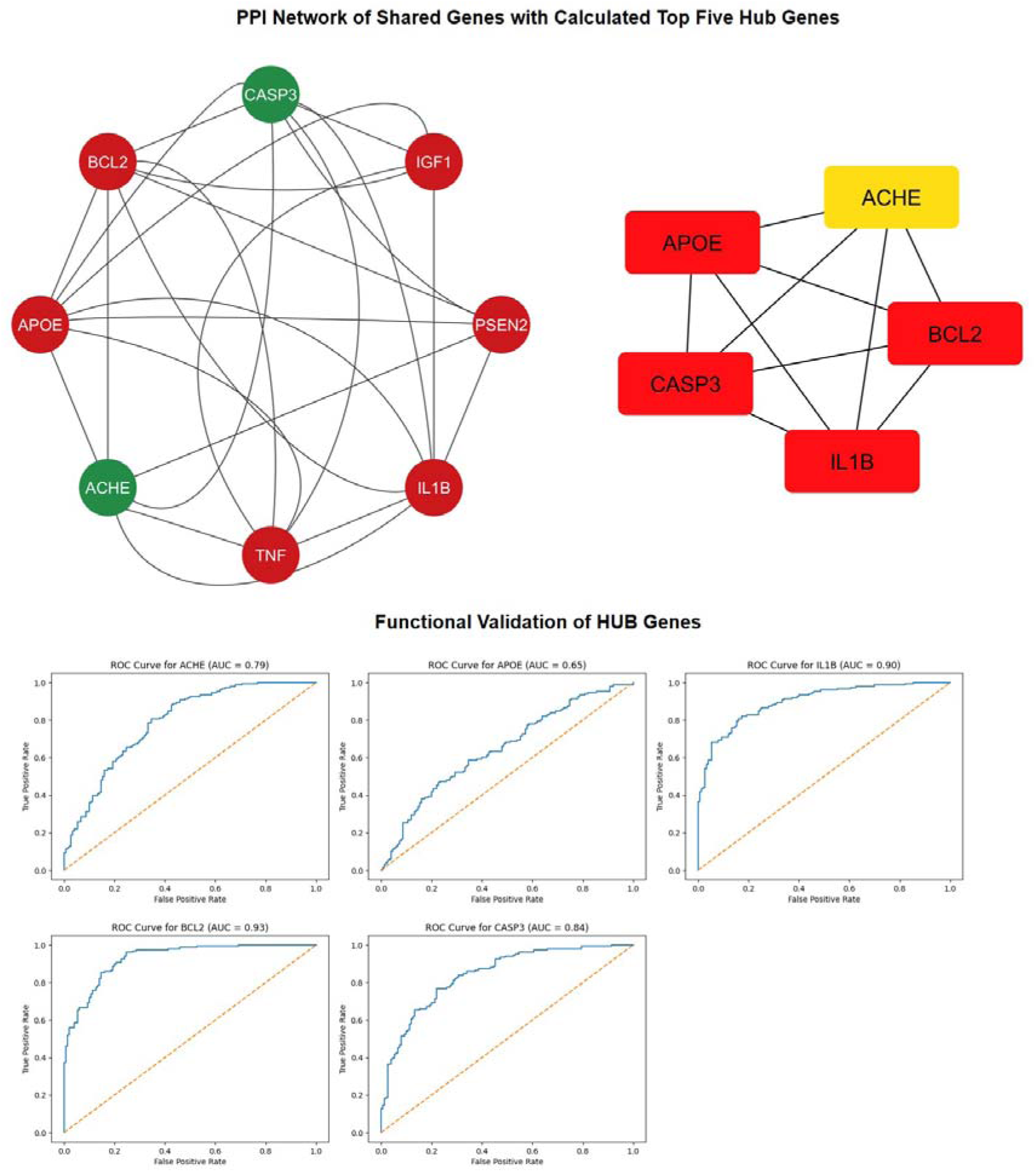
Protein-protein interaction network and diagnostic validation of hub genes linking polystyrene exposure with Alzheimer’s disease. The left panel shows the STRING-derived interaction network of genes shared between polystyrene exposure and Alzheimer’s disease. The right panel highlights the top five hub genes identified by CytoHubba topological analysis: APOE, ACHE, CASP3, IL1B, and BCL2. Lower panels show receiver operating characteristic (ROC) curve analysis of these hub genes in the GSE48350 validation dataset. BCL2 exhibited the highest diagnostic performance (AUC = 0.93), followed by IL1B (AUC = 0.90), CASP3 (AUC = 0.84), ACHE (AUC = 0.79), and APOE (AUC = 0.65).

Additionally, ACHE showed moderate predictive accuracy (AUC = 0.788), while APOE demonstrated modest but significant predictive value (AUC = 0.646) **(Table 2)**. These results indicate that these genes may serve as potential biomarkers for AD diagnosis and disease progression monitoring.

**Table 2.** Diagnostic performance of hub genes identified from the protein–protein interaction network in distinguishing Alzheimer’s disease samples from controls in the GSE48350 validation dataset. The table presents the area under the ROC curve (AUC) and corresponding 95% confidence intervals for APOE, ACHE, CASP3, IL1B, and BCL2.

| Gene | AUC | 95% CI Lower | 95% CI Upper |
| --- | --- | --- | --- |
| APOE | 0.646 | 0.585 | 0.707 |
| ACHE | 0.788 | 0.735 | 0.837 |
| CASP3 | 0.842 | 0.795 | 0.884 |
| IL1B | 0.898 | 0.86 | 0.93 |
| BCL2 | 0.929 | 0.9 | 0.954 |

## 4. Discussion

The present study applied an integrative bioinformatics framework to investigate potential molecular connections between polystyrene exposure and AD by integrating gene-disease association databases, transcriptomic analysis, WGCNA, pathway enrichment analysis, and PPI network modeling. This multilayered analytical strategy enabled the identification of 16 intersecting genes shared between AD-associated genes and polystyrene-responsive genes, several of which exhibited significant transcriptional alterations in hippocampal tissues from AD patients (Wang et al., 2024; Gaspar et al., 2025; Liu et al., 2026). Network topology analysis further revealed IL1B, CASP3, BCL2, ACHE, and APOE as central hub genes within the interaction network, suggesting that these molecules may function as critical regulators linking environmental plastic exposure with neurodegenerative signaling pathways. Accumulating evidence indicates that microplastics and nanoplastics derived from environmental polymers, such as polystyrene can accumulate in biological tissues and induce multiple cellular stress responses, including oxidative stress, inflammatory activation, and mitochondrial dysfunction. These biological effects are particularly relevant to NDDs, including AD, which is characterized by progressive neuronal loss, chronic neuroinflammation, and impaired metabolic homeostasis (Chen et al., 2025; Lafram et al., 2025b; Yu et al., 2025a). Consistent with this notion, the integrative analysis conducted in this study identified several genes associated with oxidative stress responses (SOD2, HMOX1), apoptotic regulation (BAX, BCL2, CASP3), inflammatory signaling (IL1B, TNF), and neuronal metabolic processes (IGF1, DHCR24). These pathways are widely recognized contributors to AD pathogenesis and may represent potential mechanisms through which environmental plastic exposure exacerbates neurodegenerative processes. Oxidative stress is widely considered one of the earliest pathological events in AD development (Zhang et al., 2024b). The observed upregulation of SOD2 and HMOX1, both key antioxidant response genes, suggests the activation of compensatory cellular defense mechanisms against oxidative damage in AD hippocampal tissues.

Previous studies have demonstrated that exposure to microplastics can induce excessive production of ROS and disrupt mitochondrial function, thereby promoting neuronal injury and cellular dysfunction (Bhattacharyya et al., 2025). Consequently, the dysregulation of oxidative stress-related genes identified in this analysis may reflect a biological response to environmental stressors that converge on established AD-associated neurodegenerative pathways. In addition to oxidative stress mechanisms, the present analysis highlights the importance of neuroinflammatory signaling in the molecular interplay between polystyrene exposure and AD pathology. In particular, IL1B and TNF emerged as key nodes within the AD-polystyrene interaction network. These genes encode pro-inflammatory cytokines that are essential mediators of microglial activation and neuroimmune signaling, both of which are prominent pathological features of AD (Shaftel et al., 2008; Wang et al., 2025). Chronic neuroinflammation has been shown to accelerate neuronal degeneration, enhance amyloid-beta accumulation, and impair synaptic plasticity. Supporting this observation, pathway enrichment analysis revealed significant activation of immune-related signaling pathways, including interleukin signaling, inflammasome activation, and pyroptosis-related processes, which are increasingly recognized as central contributors to NDDs progression (McGroarty et al., 2025; Tang, 2025). Further, environmental pollutants, including microplastics, have been shown to activate inflammatory pathways within the central nervous system, and there is growing evidence that such particles can penetrate biological barriers, including the BBB, thereby initiating inflammatory cascades in neural tissues (Xie et al., 2024; Shi et al., 2025). The involvement of IL1B-centered inflammatory signaling therefore suggests that environmental plastic exposure may contribute to AD progression through persistent neuroimmune activation and inflammatory dysregulation. Beyond inflammatory responses, the analysis also identified multiple genes involved in apoptotic signaling and mitochondrial dysfunction, including CASP3, BAX, and BCL2, which together form a regulatory network governing programmed cell death. The equilibrium between pro-apoptotic and anti-apoptotic signaling pathways is essential for maintaining neuronal viability, and disruption of this balance has been extensively documented in AD (Jain et al., 2024; Martinez-Perez et al., 2025). The increased expression of CASP3 and BAX, combined with the reduced expression of BCL2, suggests a shift toward pro-apoptotic signaling within AD hippocampal tissues. Activation of caspase-3 represents a pivotal step in neuronal apoptosis and has been implicated in both amyloid-beta-induced neurotoxicity and tau-mediated neurodegeneration (Harada & Sugimoto, 1999; He et al., 2018; Wójcik et al., 2024). Additionally, environmental stressors, including microplastic exposure, have also been reported to activate mitochondrial apoptotic pathways through oxidative damage and metabolic dysregulation. Therefore, the dysregulation of these apoptosis-related genes may represent a converging molecular mechanism through which environmental exposures amplify neuronal loss during AD progression (Ding et al., 2024; Gecegelen et al., 2025; Kovacs et al., 2025; Marycleopha et al., 2026). The present study also highlights the importance of lipid metabolism and synaptic signaling pathways in the molecular network linking polystyrene exposure with AD. The identification of APOE and ACHE as central hub genes within the PPI network is particularly noteworthy. APOE is widely recognized as the most significant genetic risk factor for late-onset AD and plays a fundamental role in lipid transport, neuronal repair mechanisms, and amyloid-beta clearance (Belaidi et al., 2025; Islam et al., 2025). Dysregulation of APOE expression may therefore contribute to impaired lipid metabolism and neuronal homeostasis during AD progression (Shen et al., 2025). Similarly, ACHE, which encodes acetylcholinesterase, plays a critical role in regulating cholinergic neurotransmission by hydrolyzing acetylcholine at synaptic junctions. According to the cholinergic hypothesis of AD, degeneration of cholinergic neurons contributes significantly to cognitive impairment observed in AD patients (Chen et al., 2022; Huang et al., 2022). Thus, the identification of ACHE as a hub gene within the interaction network suggests that disturbances in cholinergic signaling may represent an additional pathway through which environmental exposures influence neurodegenerative processes. In line with this interpretation, pathway enrichment analysis revealed significant involvement of synaptic signaling pathways, including GABAergic synapse, glutamatergic synapse, and synaptic vesicle cycle pathways, which are essential for neuronal communication and synaptic plasticity. Dysfunction of these pathways is widely recognized as a central feature of AD-related cognitive decline (Rivera et al., 2023). In addition, the WGCNA-based network analysis further emphasized the importance of transcriptional network organization in AD pathogenesis. The identification of dark green and light-yellow co-expression modules strongly correlated with AD clinical traits indicates that AD-related molecular alterations occur within coordinated gene networks rather than isolated gene changes. Integration of the intersecting AD-polystyrene genes with these modules revealed that several key genes, including IL1B, CASP3, APOE, and BCL2, are embedded within AD-associated co-expression networks, reinforcing their biological relevance. The strong correlations observed between module membership and gene significance further suggest that these genes occupy central positions within the transcriptional architecture of AD, highlighting their potential value as therapeutic targets. An additional important aspect of this study is the diagnostic evaluation of hub genes, which demonstrated that several genes, particularly BCL2, IL1B, and CASP3, exhibit strong predictive performance in distinguishing AD samples from healthy controls. These findings suggest that these genes may serve as potential molecular biomarkers for AD diagnosis and disease progression monitoring (Sun et al., 2019; Tsumagari et al., 2022). Given the increasing global concern regarding environmental microplastic pollution and its potential health impacts, the identification of molecular pathways linking polystyrene exposure with neurodegenerative signaling pathways has important public health implications (Chulkov et al., 2025; Zhang et al., 2025). The results of this study indicate that environmental plastic exposure may influence AD development through multifactorial mechanisms involving oxidative stress, neuroinflammation, apoptotic signaling, and synaptic dysfunction, thereby providing a novel perspective on environmental contributors to NDDs. Thus, the identification of central hub genes, including IL1B, CASP3, BCL2, ACHE, and APOE, highlights key regulatory pathways involved in neuroinflammation, oxidative stress, apoptosis, and neuronal metabolic regulation. These findings expand our understanding of the potential role of environmental plastic exposure in NDDs development and provide a foundation for future experimental studies aimed at elucidating the environmental determinants of AD.

## 5. Conclusion and Future Directions

This study employed an integrative systems biology approach to investigate potential molecular links between polystyrene exposure and AD by combining gene-disease association databases, transcriptomic analysis, WGCNA, functional enrichment analysis, and PPI network modeling. The analysis identified 16 intersecting genes shared between AD-related genes and polystyrene-associated genes, indicating a potential molecular interface between environmental plastic exposure and AD. Transcriptomic profiling further revealed significant differential expression of several of these genes in AD hippocampal tissues, suggesting their involvement in biological processes closely associated with AD pathology. Functional enrichment and network analyses indicated that these genes participate in key pathways related to neuroinflammatory signaling, oxidative stress responses, apoptotic regulation, lipid metabolism, and synaptic signaling, which represent fundamental mechanisms underlying AD progression. Notably, IL1B, CASP3, BCL2, ACHE, and APOE were identified as central hub genes within the interaction network and demonstrated promising diagnostic performance in ROC analysis, highlighting their potential value as biomarkers or therapeutic targets. Collectively, these findings provide new insights into how environmental microplastic exposure, particularly polystyrene, may contribute to AD pathogenesis through interconnected molecular mechanisms. However, several limitations should be considered. The findings are primarily based on computational analyses of publicly available transcriptomic datasets and therefore require experimental validation to confirm the biological relevance of the identified genes and pathways. In addition, the transcriptomic data analyzed in this study were derived from hippocampal tissue, which represents only one of several brain regions affected during AD progression. Furthermore, human exposure to microplastics typically involves a mixture of polymers and plastic-associated chemical additives, which may exert combined or synergistic effects that were not fully captured in this analysis. Future research should therefore focus on experimental validation using cellular and animal models of microplastic exposure, including neuronal cultures, brain organoids, and *in vivo* models, to elucidate the direct neurotoxic effects of polystyrene micro- and nanoplastics. Integrating multi-omics approaches, such as proteomics, metabolomics, and epigenomics, together with single-cell and spatial transcriptomics, could provide deeper insights into the cell-type-specific molecular mechanisms underlying microplastic-induced neurodegeneration. Additionally, epidemiological studies investigating associations between environmental microplastic exposure and cognitive decline in human populations will be essential for understanding the broader public health implications of environmental plastic pollution in relation to AD.

## Data Availability Statement

The original contributions presented in the study are included in the article/Supplementary Material, further inquiries can be directed to the corresponding author/s.

## Authors’ Contribution

**Rohan Gupta**: Data curation, Writing- Original draft preparation, Formal analysis, Visualization, Conceptualization, Visualization, Investigation, Formal analysis, Writing- Reviewing and Editing; **Sorabh Lakhanpal:** Writing- Original draft preparation, Formal analysis, Visualization; **Saurabh Gupta:** Writing- Original draft preparation, Formal analysis, Visualization; **Niraj Kumar Jha**: Formal analysis, Writing- Reviewing and Editing; Visualization;

## Acknowledgement

We would like to thank the senior management of Galgotias University for their support.

## Supplementary Materials

The Supplementary Material for this article can be found online.

## Gels and Blots Image(s)

Not Applicable

