## Supplementary material for "A Systems Neuroscience Approach Identifies IL1B-CASP3 Signaling as a Molecular Link Between Polystyrene Exposure and Alzheimer’s Disease": Supplemnetary Data: Supplementary Figures.docx

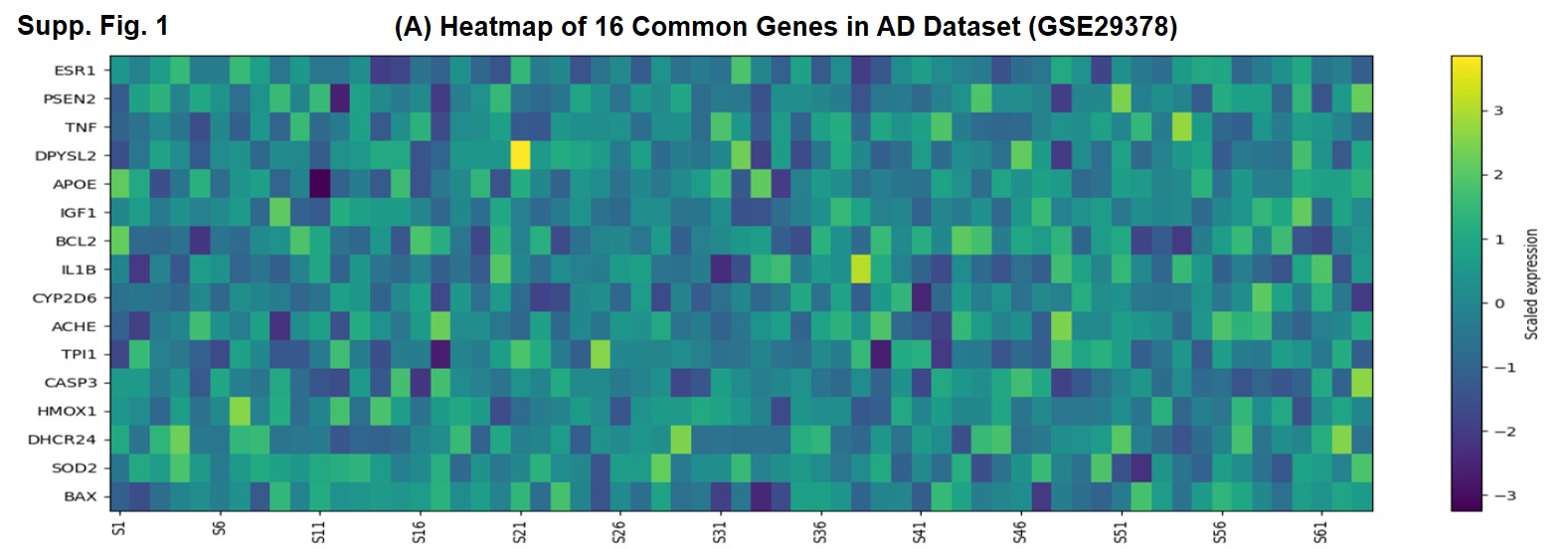

**Supplementary Figure 1: Heatmap showing expression profiles of the 16 intersecting genes shared between polystyrene exposure and Alzheimer's disease in the GSE29378 hippocampal dataset. Rows represent candidate genes and columns represent individual samples. Color intensity corresponds to scaled gene expression values, with yellow indicating higher expression and purple indicating lower expression. The heatmap illustrates heterogeneous but distinct expression patterns of genes involved in neuroinflammation, oxidative stress, apoptosis, and neuronal signaling across Alzheimer’s disease samples.**

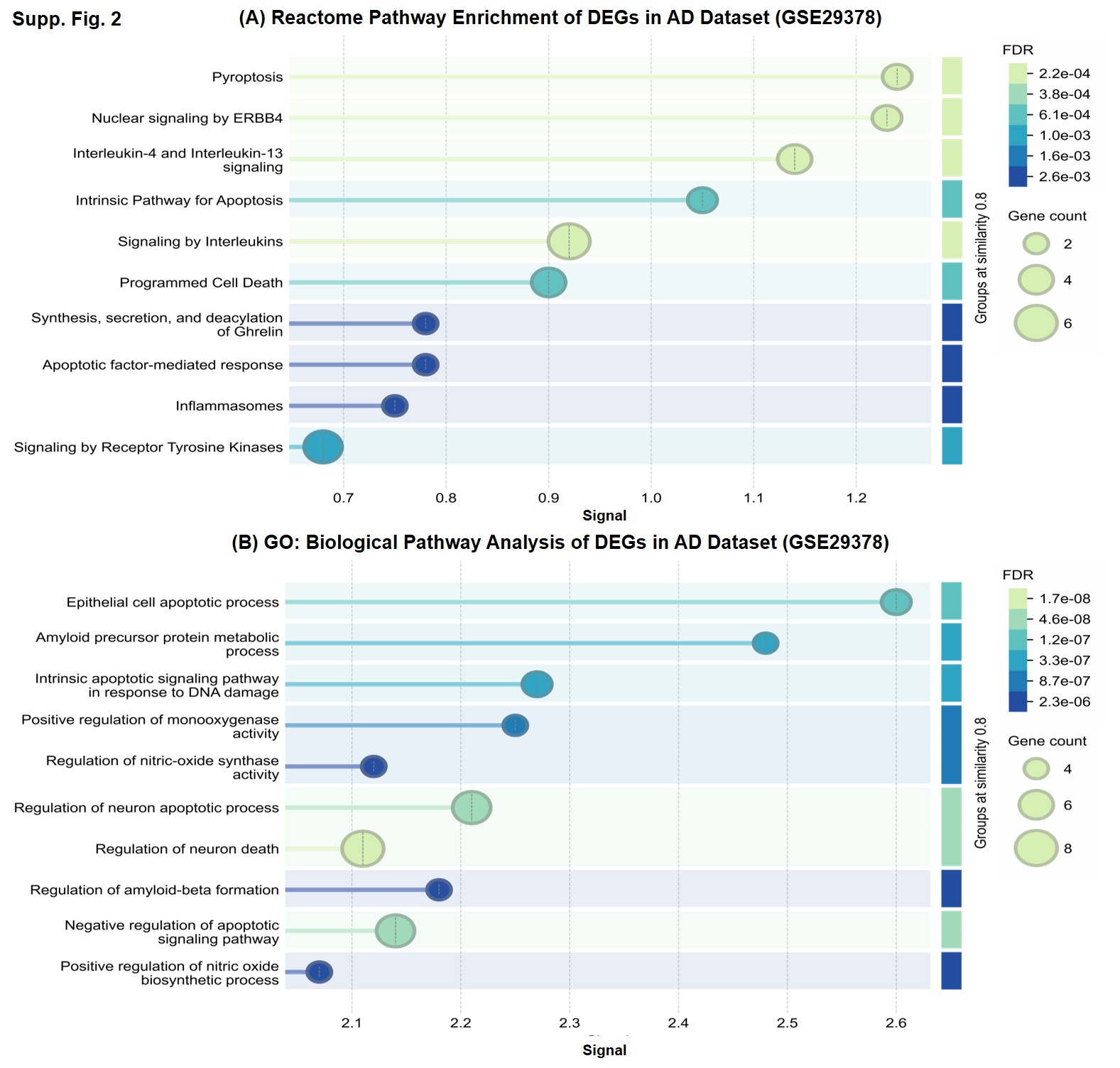

**Supplementary Figure 2: Functional enrichment analysis of differentially expressed genes in the GSE29378 hippocampal dataset from Alzheimer's disease patients. (A) Reactome pathway enrichment analysis showing significant enrichment of pathways related to pyroptosis, interleukin signaling, apoptosis, programmed cell death, inflammasome activation, receptor tyrosine kinase signaling, and ERBB4 signaling. (B) Gene Ontology biological process enrichment analysis showing significant enrichment of pathways involved in epithelial and neuronal apoptosis, amyloid precursor protein metabolism, amyloid-beta formation, nitric oxide biosynthesis, and regulation of neuron death. Bubble size represents the number of genes enriched in each pathway, whereas color indicates the false discovery rate (FDR).**

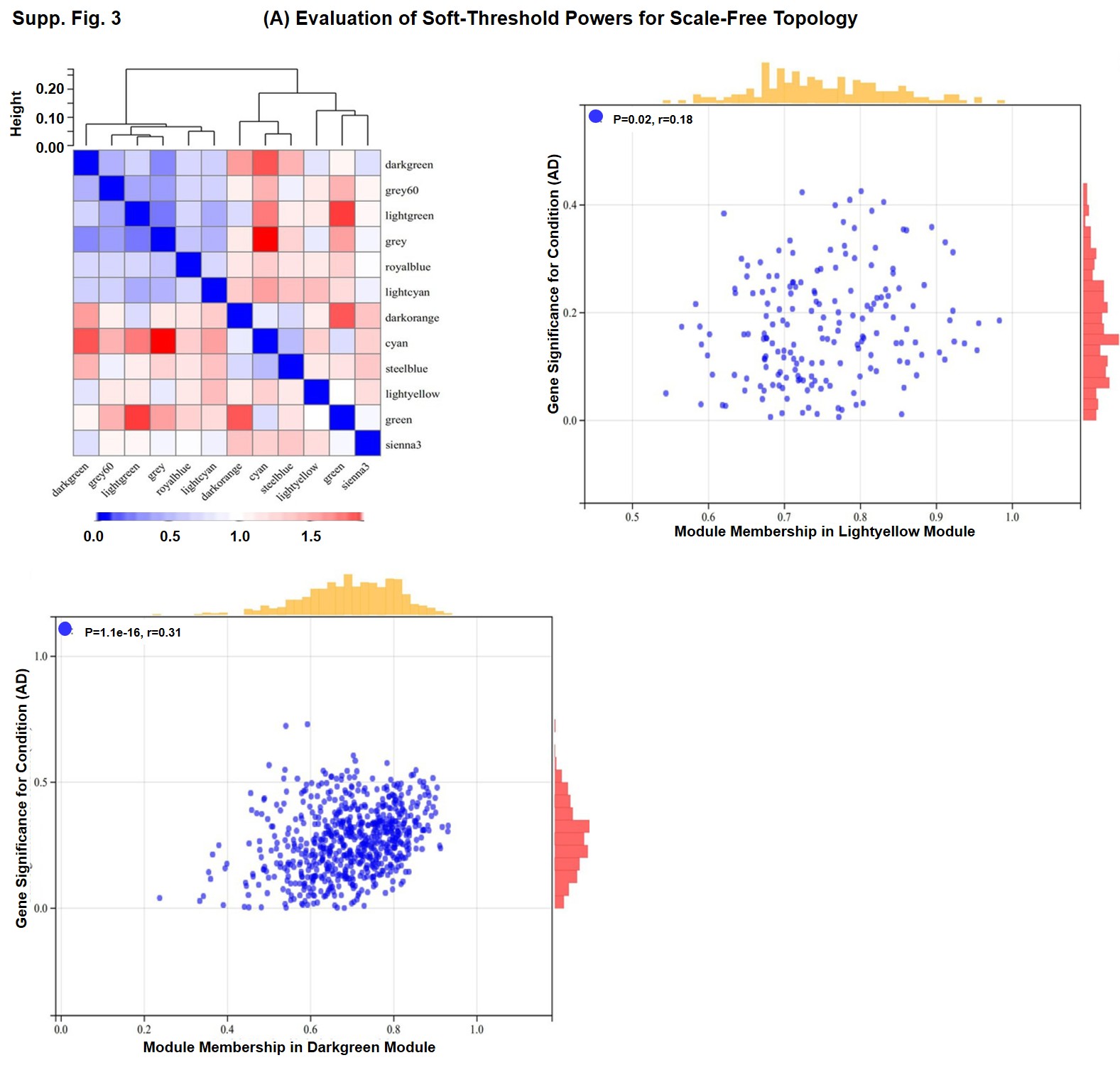

**Supplementary Figure 3: Correlation analysis of WGCNA modules associated with Alzheimer's disease in the GSE29378 dataset. The upper left panel shows hierarchical clustering of module eigengenes, illustrating relationships among the identified co-expression modules. Red indicates stronger positive correlation and blue indicates weaker correlation between module eigengenes. The upper right and lower panels show scatter plots of module membership versus gene significance for AD in the light-yellow and dark green modules, respectively. Positive correlations between module membership and gene significance indicate that highly connected genes within these modules are strongly associated with Alzheimer’s disease status.**
